# Mitochondrial fusion mediated by mitofusin 1 regulates macrophage mycobactericidal activity by enhancing autophagy

**DOI:** 10.1101/2020.12.01.407544

**Authors:** Yuping Ning, Yi Cai, Youchao Dai, Siwei Mo, Oliver Werz, Xinchun Chen

**Author notes:** Address correspondence to Xinchun Chen, or Oliver Werz,. Y.N and Y.C contributed equally to this work.

## Abstract

Mitochondria as a highly dynamic organelle continuously changes morphology and position during its life cycle. Mitochondrial dynamics including fission and fusion play a critical role in maintaining functional mitochondria for ATP production, which is directly linked to host defense against Mtb infection. However, how macrophages regulate mitochondrial dynamics during *Mycobacterium tuberculosis* (Mtb) infection remains elusive. In this study, we found that Mtb infection induced mitochondrial fusion through enhancing the expression of mitofusin 1 (*MFN1*), which resulted in increased ATP production. Silencing *MFN1* inhibited mitochondrial fusion and subsequently reduced ATP production, which, in turn, severely impaired macrophage mycobactericidal activity by inhibiting autophagy. Impairment of mycobactericidal activity and autophagy was replicated using oligomycin, an inhibitor of ATP synthase. In summary, our study revealed MFN1-mediated mitochondrial fusion is essential for macrophage mycobactericidal activity through the regulation of ATP dependent autophagy. MFN1-mediated metabolism pathway might be targets for development of host direct therapy (HDT) strategy against TB.

**Importance:** How mitochondrial dynamic is regulated in attempt to fight against Mtb remains elusive. Our study revealed that the fission/fusion dynamics of mitochondria during Mtb infection is regulated by MFN1, through which mitochondrial respiration and autophagy activity are affected. Our findings suggested that intervention of energy metabolism by targeting MFN1 might be a strategy against Mtb infection.

## Introduction

Tuberculosis (TB), caused by intracellular infection with the bacteria *Mycobacterium tuberculosis* (Mtb), is a leading infectious disease worldwide that killed proximately 1.4 million people in 2018 (1). Macrophages are the front-line responders to pathogens, act as either reservoirs or killer of the bacteria (2). Immunometabolism, which supports the function and bactericidal activity of macrophages, has recently been attracting a lot of interest (3). To eliminate Mtb efficiently, macrophages have been reported shift their metabolic states to glycolysis and oxidative phosphorylation (OXPHOS) to meet the bioenergetic and metabolic requirements (4).

Mitochondria are highly dynamic powerhouses that control cell energy metabolism (5). They are known to efficiently produce the bulk of the ATP needed to support cellular activities through OXPHOS and the TCA cycle (6, 7). Mitochondrial dynamics, including fission and fusion, primarily regulated by mitofusins (MFN) and dynamin-related protein 1 (Drp1), are related to the metabolic processes of OXPHOS and glycolysis (8, 9). Mitochondrial fusion increases dimerization and the activity of ATP synthase, maximizes the oxidative capacity, and maintains mitochondrial function (10).

During Mtb infection, biphasic immunometabolism, which involves changes in mitochondrial function, occurs in macrophages (11). However, how Mtb infection influences mitochondrial dynamics and the metabolic profile remains uncertain. A previous study found that different Mtb virulent factors altered differential mitochondrial dynamics in macrophages (12). Jamwal et al. (2012) showed that, compared with an avirulent strain, a virulent strain of Mtb elongated the mitochondria and increased ATP levels in a human macrophage THP-1 cell line as a way to prevent host macrophage apoptosis (13). In contrast, Coulson et al. (2015) and Lee et al. (2019) found that Mtb induced mitochondrial fission in a human cell line A549 (14) and murine cell line BMDM (15), respectively.

In this study, we found that mitochondria undergo fusion during Mtb infection mediated by MFN1, leading to significantly increased OXPHOS levels and ATP production. Silencing *MFN1* inhibited mitochondrial fusion and subsequently reduced ATP production, which, in turn, severely impaired macrophage mycobactericidal activity by inhibiting autophagy.

## Results

### Mtb infection induces mitochondrial fusion in human macrophages

To investigate the effects of Mtb infection on mitochondrial dynamics, we infected THP-1 macrophages with the Mtb strains H37Ra and H37Rv and monitored the mitochondrial morphology using confocal microscopy. We found that Mtb-H37Ra-infected macrophages displayed a higher rate of mitochondrial fusion and more tubular mitochondrial networks than uninfected macrophages (Fig. 1 A, B and Fig. S1 A). The same phenomenon was observed with the virulent Mtb strain, H37Rv (Fig. 1 C, D). Mitochondrial fusion has been associated with increased mitochondrial membrane potential (MMP) (16, 17). We thus further investigated the MMP during Mtb infection using MitoTraker. The MMP was significantly elevated in both H37Ra- and H37Rv-infected THP-1 macrophages at 24 to 48 hours postinfection (hpi) (Fig. 1 E, F). These results were further confirmed by using another MMP dye, rhodamine 123 (18). We observed a decreased intensity of rhodamine 123 staining in Mtb-infected macrophages, indicative of increased MMP (Fig. S1 B). Together, these results indicated that mitochondrial dynamics were altered by Mtb infection, and the mitochondria tended to form reticulate, elongated fusion networks.

**Figure 1.**
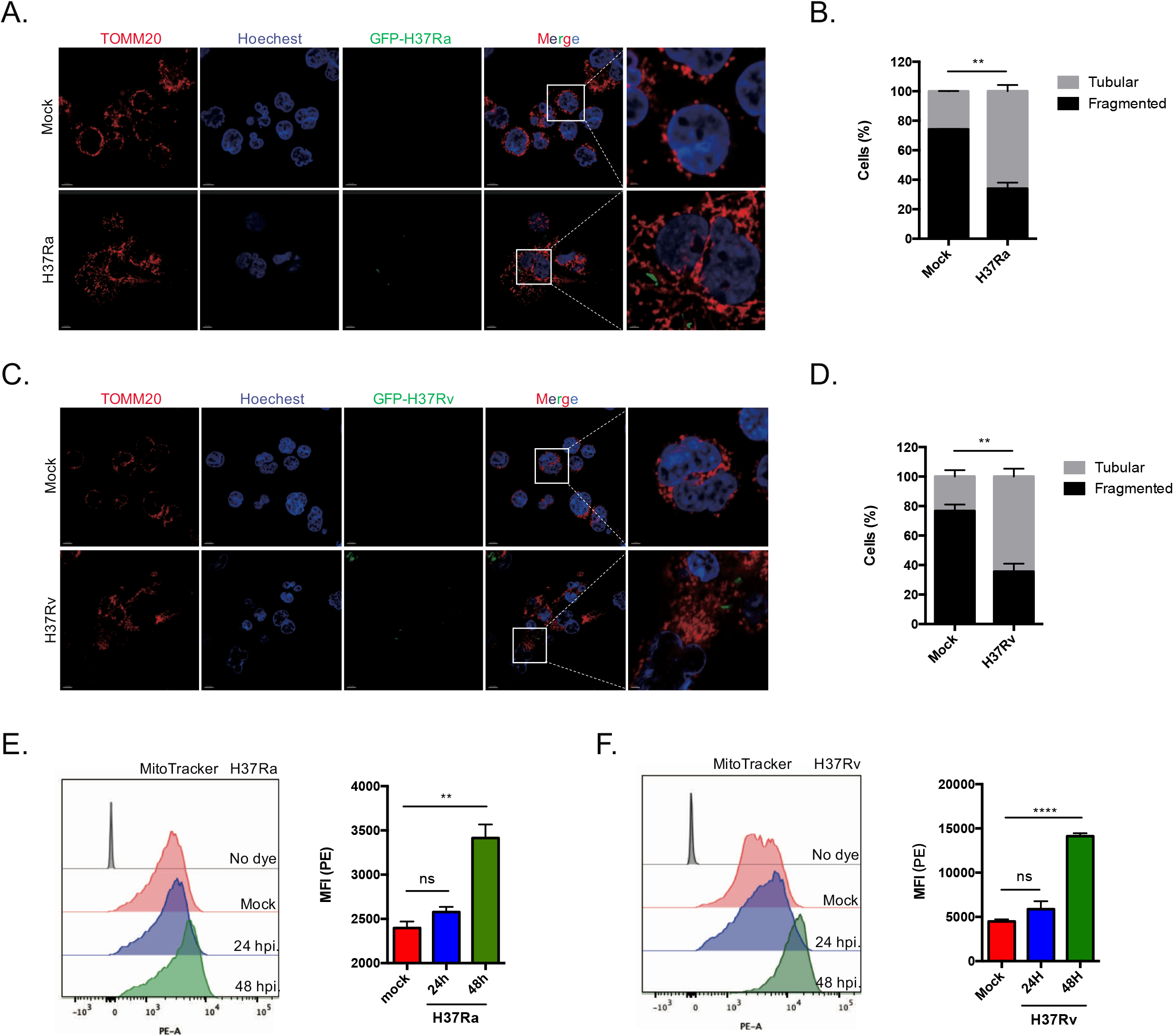
Mitochondrial fusion during Mtb infection. A-D. Confocal microscopy images showing elongated mitochondrial morphology in THP-1 macrophages infected by H37Ra (MOI = 10:1) and H37Rv (MOI = 10:1) for 48 h. Red fluorescence, mitochondria; green fluorescence, H37Ra/Rv; blue fluorescence, Hoechst-33342-labeled nuclei. (Scale bars, left 10 μm, right 3 μm) (A, C) and histograms (n = 200 cells) (B, D) of the percentage of macrophages with “fragmented” mitochondria compared with those with “tubular” mitochondrial morphologies in the mock and H37Ra/Rv-infected groups. E-F. Mitochondrial membrane potential (MMP) was detected using MitoTracker Red CMXRos probs (100 nM) in H37Ra- (MOI = 10:1) (E) and H37Rv- (MOI = 10:1) (F) infected THP-1 macrophages. Mean fluorescence intensity (MFI) of MMP was analyzed by flow cytometry. Histograms indicated MFI in the mock and H37Ra/Rv-infected groups. The data represent the means ± SEM (n = 3). **, p < 0.01; ****, p < 0.0001; “ns”, not significant.

### Mtb-induced mitochondrial fusion depends on MFN1

Key players in the mitochondrial fusion process include the outer mitochondrial membrane GTPases mitofusin 1 (MFN1) and mitofusin 2 (MFN2), and the inner membrane GTPase optic atrophy 1 (OPA1) (19). To determine whether those proteins are involved in Mtb-mediated mitochondrial fusion, we analyzed the expression of these genes in Mtb-infected macrophages.

The results show that the expression of *MFN1* significantly increased (P<0.0001) in both H37Ra- and H37Rv-infected macrophages (Fig. 2 A, B). The level of *MFN1* protein was consistently found to be significantly higher in the Mtb-infected macrophages than the uninfected cells (Fig. 2 C, D). On the contrast, the levels of MFN2 and OPA1 are not significantly altered. Notably, elevated MFN1 level was also observed in H37Ra-infected primary human monocyte-derived macrophages (hMDMs) (Fig. S2). Considering the association between MFN1 and mitochondrial fusion, we therefore further investigate the role of MFN1 in regulating mitochondrial fusion in human macrophage induced by Mtb infection. To this end we knocked down *MFN1* with siRNA (Fig. 2 E) and evaluated its effect on the mitochondrial fusion. As expected, we found that silence of MFN1 dramatically reduced mitochondrial fusion during Mtb infection (Fig. 2 F, G). A significant reduction in MMP in the *MFN1*-knockdown macrophages was observed by using MitoTracker staining (Fig. 2 H). The effect of silencing MFN1 were further confirmed by JC-1 staining using confocal microscopy (Fig. S3). Compared with si*MFN1* group, control group show greater red fluorescence represent JC-1 monomers which show green florescence fuse into mitochondrial matrix and form red aggregates due to higher MMP (20). Together, these findings indicate that Mtb infection induces mitochondrial fusion by increasing *MFN1* expression.

**Figure 2.**
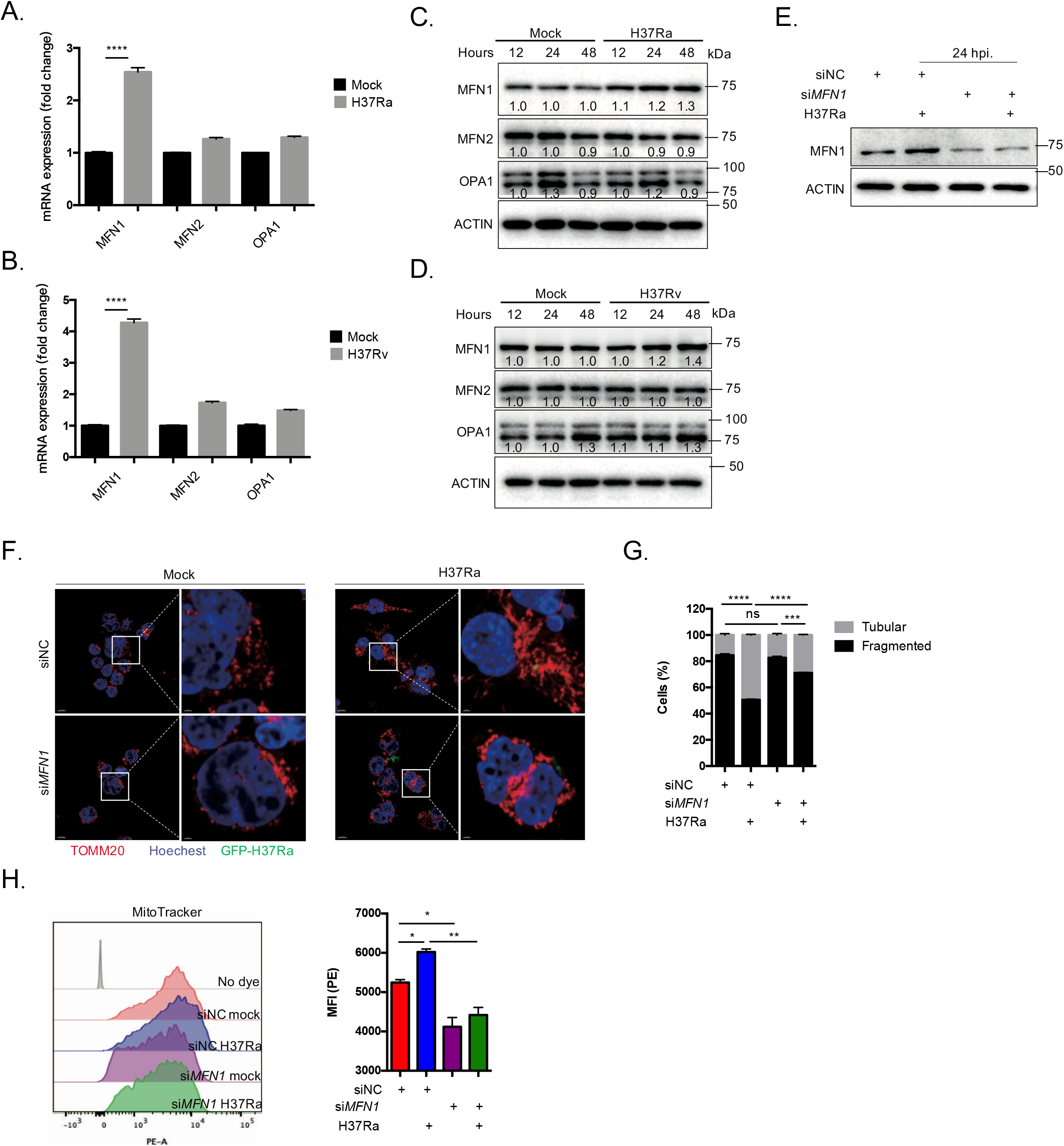
Mtb-induced mitochondrial fusion depends on MFN1. A-D. THP-1 macrophages infected with H37Ra (MOI = 10:1) or H37Rv (MOI = 10:1); the mRNA (A, B) and protein (C, D) expression of mitochondrial-fusion-related genes were analyzed by real-time-PCR and western blotting (WB), respectively. E. WB analysis of siRNA-mediated *MFN1* knockdown in THP-1 macrophages. F-G. Confocal microscopy images (F) showing mitochondrial morphology in THP-1 macrophages infected by H37Ra (MOI = 10:1) for 48 h with the negative control siRNA (siNC) and MFN1 siRNA (si*MFN1*) treatment. Red fluorescence, mitochondria; green fluorescence, H37Ra; blue fluorescence, Hoechst 33342-labeled nuclei (Scale bars, left 10 μm, right 2 μm). and histograms (n = 200 cells) (G) showing percentage of macrophages with “fragmented” mitochondria compared with those with “tubular” mitochondria. H. Mitochondrial membrane potential was detected using MitoTracker Red CMXRos probs (100 nM) in H37Ra (MOI = 10: 1) infected THP-1 macrophages 48 hpi. The data represent mean ± SEM (n = 3). *, p < 0.05; **, p < 0.01; ***, p < 0.001; ****, p<0.0001; “ns”, not significant.

### Silencing *MFN1* impairs OXPHOS and ATP production during Mtb infection

Increasing mitochondrial fusion indicates a heightened requirement for oxidative phosphorylation (OXPHOS) and ATP production (21). In line with this, we observed that Mtb infection increased ATP production and the oxygen consumption rate (OCR) (Fig. 3 A and Fig. S4) at 24 hpi. Given the role of MFN1 in Mtb-induced mitochondrial fusion, we investigated whether MFN1 expression alters the respiratory capacity during Mtb infection. As expected, the silencing of *MFN1* resulted in a dramatically decreased basal OCR and maximum OCR in the Mtb-infected macrophages (Fig. 3 B). A decrease of ATP production was also observed in the *MFN1*-knockdown macrophages during Mtb infection (Fig. 3 C). Silencing *MFN1* expression had almost no effect on mROS production of Mtb-infected and uninfected cells (Fig. S5). On the contrast, no significant changes in the extracellular acidification rate (ECAR) were found between the control group and the *MFN1* knockdown group (Fig. 3 D), suggesting that the increase in intracellular ATP production induced by Mtb was via OXPHOS. These results indicate that silencing *MFN1* impairs OXPHOS and ATP production during Mtb infection.

**Figure 3.**
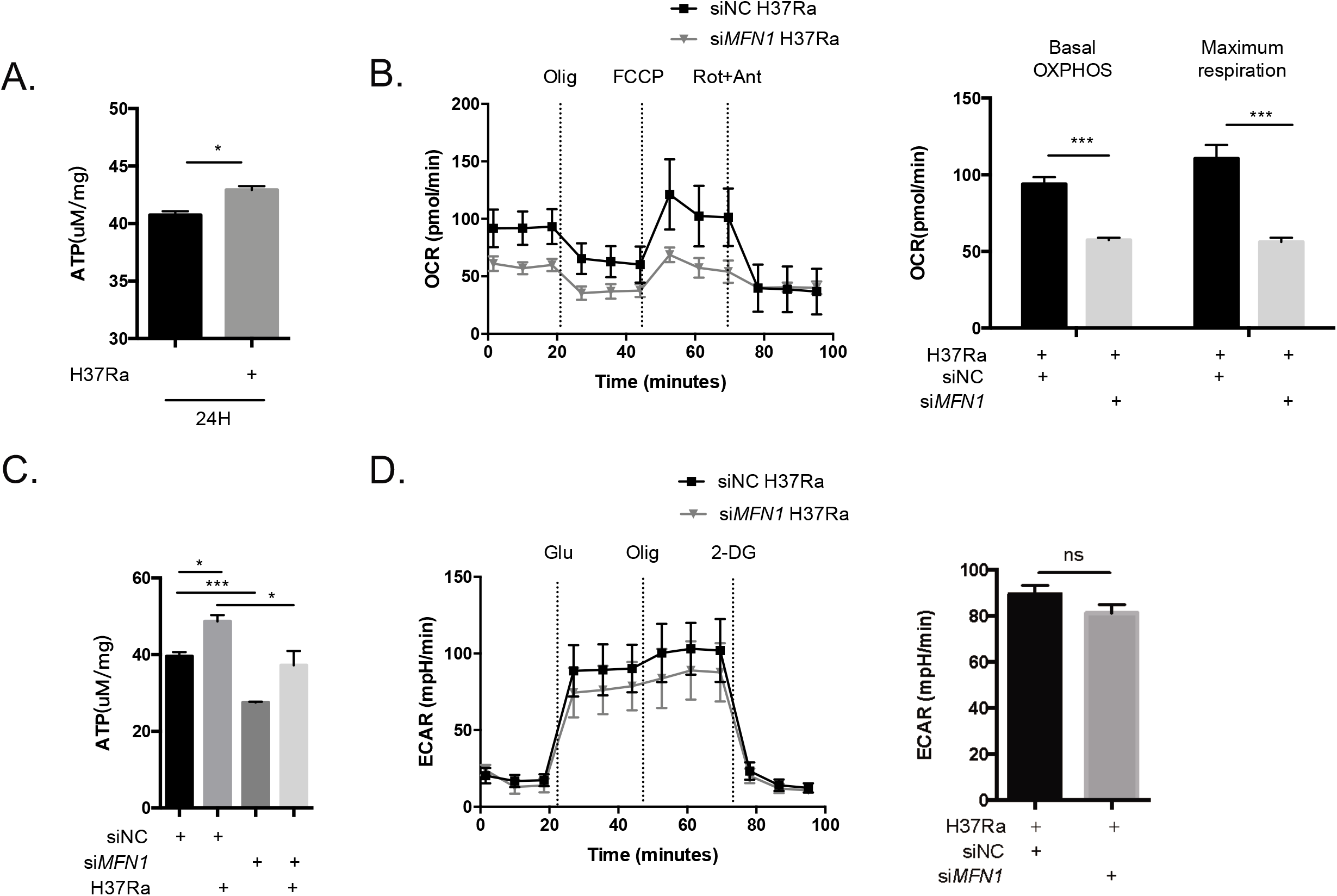
Silencing *MFN1* expression impairs Mtb-induced OXPHOS and ATP production in THP-1 macrophages. A. THP-1 macrophages were infected with H37Ra as indicated at MOI = 10:1, and total intracellular ATP was measured at 24 hpi using luminescent assay. B. Mito Stress assay detected by Seahorse XFe24 Extracellular Flux Analyzer showing change in OCR of THP-1 macrophages after infection with H37Ra (MOI = 10:1) for 24 h. The histogram on the right shows decreased basal OCR and maximum OCR in si*MFN1* cells compared with negative control siRNA (siNC) cells infected with H37Ra. C. THP-1 macrophages were treated with siNC or si*MFN1* and then infected with H37Ra (MOI = 10:1) for 24 h; the intracellular ATP level was assessed using luminescent assay. D. Glycolysis stress assay showing change in ECAR of THP-1 macrophages after infection with H37Ra (MOI = 10:1) for 24 h in the presence or absence of *MFN1* siRNA. The data represent mean ± SEM (n = 3). *, p < 0.05; ***, p < 0.001; “ns”, not significant.

### MFN1 regulates macrophage mycobactericidal activity depending on OXPHOS and ATP production

To explore whether MFN1-regulated mitochondrial fusion and respiratory capacity are linked to the host control of Mtb replication, we tested the effect of silence of *MFN1* on intracellular Mtb growth in THP-1 macrophage. We found that silencing *MFN1* significantly enhanced both Mtb H37Ra and H37Rv growth in THP-1 macrophages at 72 hpi (Fig. 4 A-B). Since si*MFN1* influence OXPHOS and ATP production, we propose that the impaired mycobactericidal activity observed in si*MFN1*-macrophage is due to the reduced ATP production. To test this, we applied oligomycin, an inhibitor of ATP synthase, to pharmacologically inhibit OXPHOS (22, 23) and evaluated its effect on intracellular Mtb growth within THP-1 macrophage. At a concentration of 10 nM, oligomycin dramatically decreased intracellular ATP in macrophage with or without Mtb infection (Fig. 4 C). When intracellular CFU numbers were recorded for the oligomycin treated and untreated groups at 6 and 72 hpi, we confirmed that oligomycin treatment significantly impaired the THP-1 cells mycobactericidal activity (Fig. 4 D).

**Figure 4.**
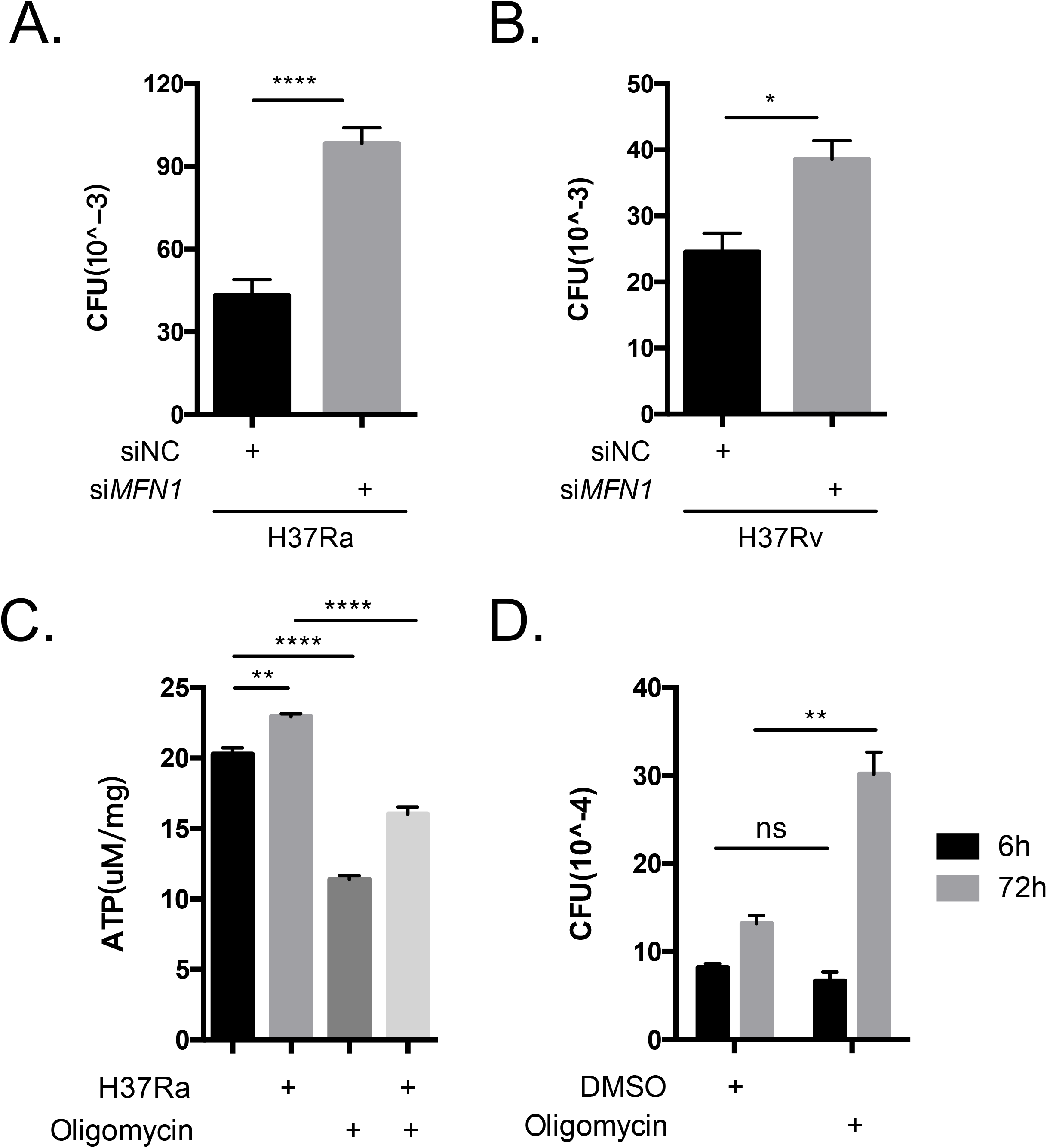
MFN1-mediated macrophage mycobactericidal activity depends on OXPHOS and ATP production. A-B. THP-1 macrophages were infected with H37Ra (MOI =10:1) (A) or H37Rv (MOI = 10:1) (B) in the present or absence of *MFN1*-siRNA compared with negative control siRNA (siNC) for 6 h, then, extracellular bacteria were removed and the intracellular CFU was determined at 72 hpi. C. THP-1 macrophages treated with oligomycin (10 nM) were infected as indicated at MOI = 10:1, and total intracellular ATP was measured at 24 hpi using luminescent assay. D. THP-1 macrophages treated with DMSO or oligomycin (10 μM) were infected with H37Ra (MOI = 10:1) and CFU counts were recorded at 6 hpi; for the 72 h CFU counting, extracellular bacteria were removed at 6 hpi, then, macrophage bactericidal activity was assessed by CFU counts of intracellular H37Ra at 72 hpi. The data represent mean ± SEM (n = 3). *, p < 0.05; **, p < 0.01; ****, p<0.0001; “ns”, not significant.

Thus, our results indicated that MFN1 expression induced by Mtb infection is a mechanism for host macrophage to get rid of intracellular Mtb by modulating mitochondrial fusion, a process with enhanced OXPHOS and ATP production.

### MFN1 enhanced macrophage bactericidal activity against Mtb by enhancing autophagy

Mitochondrial dynamics, including fusion and fission, are essential for modulating mitochondrial bioenergetics and a variety of cellular processes, including autophagy (24). Since autophagy is a main strategy for host defense against Mtb, we first investigated the effect of MFN1 on autophagy induction during Mtb infection. We found that silencing *MFN1* significantly inhibited autophagy in Mtb-infected macrophages, as evidenced by a significant decrease in LC3B-II (Fig. 5A). Secondly, we used autophagosome-lysosome fusion inhibitors, chloroquine and bafilomycin A1, which lead LC3B-II accumulation and present a higher level of LC3B-II (25), to confirm whether reduced LC3B-II signal in si*MFN1* group caused by inhibition of autophagy flux rather than acceleration of it. We found significantly decreased LC3B-II level in si*MFN1* compared with control macrophage both treated with chloroquine and bafilomycin A1 (Fig. S5 A), indicating si*MFN1* inhibit the initiation of autophagy process, rather than acidification of autophagosome. In consistent with the role of ATP we proposed in MFN-mediated mycobactericidal activity, we found that inhibition of mitochondrial respiration by oligomycin significantly reduced the level of LC3B-II (Fig. 5 B and Fig. S6 B) during Mtb infection. The link between the inhibition of cellular ATP production and autophagy were further confirmed by inhibitors of the electron transport chain (Fig. S6 C-D). Next, we then used a stably transformed mRFP-GFP-LC3 reporter in the THP-1 macrophages (26) to further determine the effect of MFN1 on autophagy flux. Silencing of *MFN1* significantly reduced the numbers of autophagosomes and autolysosomes, which formed puncta structures in H37Ra-infected cells, (Fig. 5 C-E), indicating that MFN1 is necessary for autophagy during Mtb infection.

**Figure 5.**
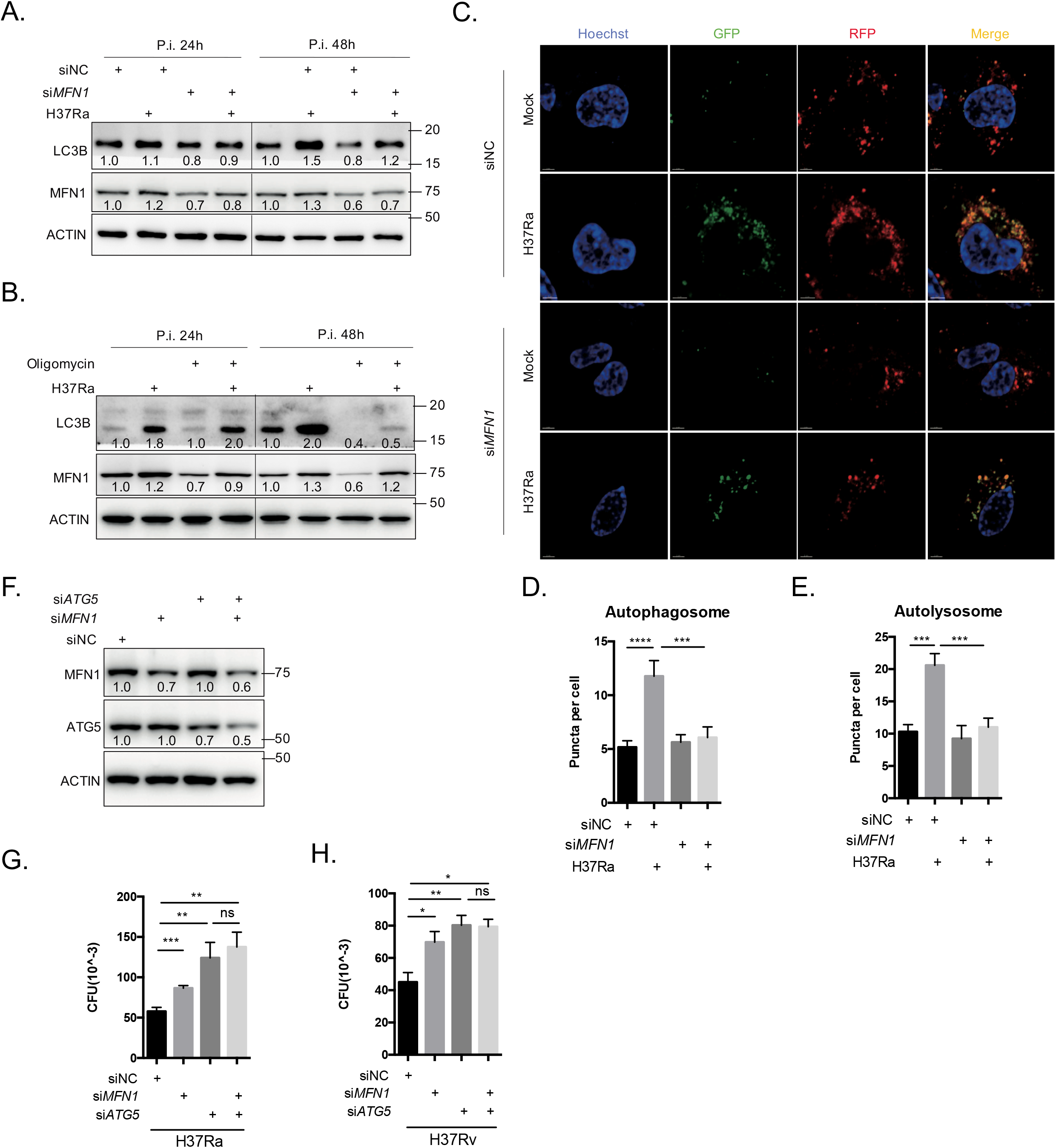
MFN1 mediated intracellular Mtb elimination by enhancing autophagy. A-B. Cell lysates were harvested from uninfected (mock) and 24-48 h H37Ra (MOI = 10: 1) infected THP-1 macrophages treated with or without *MFN1* siRNA compared with negative control siRNA (siNC) (A) and oligomycin (B). Western blot analysis of MFN1 and LC3B-II expression; β-actin was used as the loading control. C-E. Confocal microscopy images (C) and histograms (D-E) (n = 30-40 cells) showing autophagy flux in mRFP-GFP LC3 reporter THP-1 macrophages infected by H37Ra (MOI = 10: 1) for 24 h with siNC and *siMFN1* treatment. (Scale bars, 5 μm). When the newly autophagosome formed, both RFP- and GFP-LC3-II exist, show yellow florescence; when autophagosome fused with lysosome, GFP diminished, show red florescence, indicate the formation of autolysosome. F. Cell lysates were harvested from uninfected (mock) and 24 h H37Ra (MOI = 10: 1) infected THP-1 macrophages treated with and *ATG5* siRNA and *MFN1* siRNA. Western blot analysis for ATG5, MFN1; β-actin was used as the loading control. G-H. THP-1 macrophages treated with or without *MFN1* and *ATG5* siRNA were infected with H37Ra (MOI = 10:1) (G) and H37Rv (MOI = 10:1) (H) for 6 h, then extracellular bacteria were removed, intracellular CFU number was determined at 72 hpi. The data represent mean ± SEM (n = 3). *, p < 0.05; **, p < 0.01; ***, p < 0.001; ****, p < 0.0001; “ns”, not significant.

Last, we confirm the role of autophagy in MFN1-mediated suppression of intracellular Mtb growth. To this end, we knocked down *ATG5,* which encodes a key protein for autophagy initiation and processing (27), and analyzed its effect of MFN1-mediated macrophage bactericidal activity. *ATG5* expression was significantly reduced following treatment with *ATG5*-specific siRNA for 48 h (Fig. 5 F). As expected, when ATG5 is intact, silencing *MFN1* lead to higher CFU number of both intracellular H37Ra and H37Rv. However, when ATG5 is knockdown, the effects of siMFN1 on intracellular Mtb H37Ra and H37Rv replication are no longer observed. (Fig. 5 G-H). Together, these results demonstrated that the MFN1 enhanced intracellular Mtb survival by reducing autophagy.

## Discussion

The production of adequate energy is essential to fit the immunometabolic demand by macrophage to fight against Mtb infection (28). In this study, we demonstrated the mitochondrial fusion events that take place in Mtb-infected human THP-1 macrophages is regulated for host defense against Mtb. We first found that Mtb infection induced expression of MFN1, a GTPase located on the mitochondrial outer membrane, has a vital role in mitochondrial fusion. MFN1-mediated mitochondrial fusion enhances OXPHOS and ATP production. This role of MFN1 is linked to macrophage bactericidal activity, which is attribute to modulation of autophagy via mitochondrial OXPHOS and ATP production.

The importance of mitochondrial dynamics during Mtb infection has been recently recognized (12-15). However, its role in bactericidal activity and underlying mechanism remains uncertain. For example, Lee et al. reported mitochondria in murine macrophages undergo fission, involving the degradation of MFN2, during H37Ra and H37Rv infections (15). This dynamic change causing apoptosis is attributable to the defensive processes of the host cell (15). In line with this report, we also found Mfn2 is decreased in murine macrophage Raw 264.7 (Fig. S6). However, similar result is not observed in human macrophages in our study, suggesting difference of mitochondrial dynamics between human and mouse. Indeed, as reported by Vijayan et al. (2019), human macrophages rely more heavily on OXPHOS than glycolysis for ATP production when stimulated with lipopolysaccharide (LPS), which is completely contrast to murine macrophages (29).

MFN2, a homolog of MFN1, plays an important role in the innate immunity against Mtb (15, 30-32). Recently, Tur et al. (2020) found MFN2 has an important function in maintaining the autophagy process in BMDM (31); whereas, in the absence of MFN2, autophagosome accumulation is dramatically diminished (31). Lloberas et al. (2020) further suggest decreased ROS release in the absent of MFN2 contribute to impaired autophagy and apoptosis (32). This suggests there is a strong association between mitochondrial dynamics and autophagy. Despite as a high homolog of MFN2, MFN1 has distinct functions and shows higher affinity to GTP (31), which means MFN1 has greater activity to regulate mitochondrial fusion and metabolism reprogramming (21). During metabolism reprogramming, MFN1 shifts the cellular metabolism from aerobic glycolysis to OXPHOS (33) and leading to higher ATP production (21). To our knowledge, our report is the first one to link the function of MFN1 as a newly innate immune modulator.

In general, autophagy is initiated to recover ATP levels and avoid cell death when cellular energy is limited, such as during increased AMP/ATP or amino acid starvation (34). However, cellular ATP levels are also fundamental for autophagy initiation, and are required in several steps during autophagy (35, 36). Mitochondrial fusion maximizes OXPHOS (37) and increases ATP production (38), which is important for supplying the immune cell’s metabolic and other energy-dependent processes (39). Moreover, a mitochondrial respiratory deficiency and loss of MMP have also been reported to inhibit autophagy by upregulating PKA activity (40), a phenomenon with which our results seem to concur. Thus, we support the hypothesis that mitochondrial respiratory capacity and ATP production contribute to autophagy against Mtb.

In conclusion, we firstly discovered that MFN1-mediated mitochondrial fusion in human THP-1 macrophages is a defensive strategy against Mtb. Mitochondrial fusion leads to an increase in mitochondrial OXPHOS and intracellular ATP production, which fuels the autophagy process to inhibit the intracellular growth of Mtb. Therefore, host-directed therapy targeting mitochondrial dynamics and metabolism reprogramming might become a potential candidate for Mtb treatment research.

## Materials and methods

### Mtb culture and infection

Mtb H37Ra and Mtb H37Rv strains were cultured in 7H9 broth (BD Biosciences, San Jose, CA, USA) supplemented with 10% Middlebrook OADC Enrichment (BBL) containing 0.05% Tween 80 (Sigma-Aldrich) and 0.2% glycerol (Sigma-Aldrich, Merck, Darmstadt, Germany) for 1 to 2 weeks at 37°C with shaking to the mid-logarithmic phase (OD_600_ = 0.3 to 0.8) prior to experiments. In all experiments involving mycobacterial infection, THP-1 macrophages were infected with Mtb H37Ra or H37Rv at a MOI of 10 mycobacteria: 1 macrophage. Before infection, the 7H9-cultured bacteria were washed once with PBS and resuspended in serum-free RPMI 1640 media, then sonicated for 5 min to obtain a single-cell suspension.

### Cell culture, macrophage differentiation, and drug treatments

The human monocytic cell line THP-1 (ATCC) and THP-1 macrophages which transformed with an mRFP-GFP-LC3B reporter (Kindly provided by Hongbo Shen, Institute Pasteur of Shanghai, Shanghai, China) (26) were cultured in RPMI-1640 (Corning, NY, USA) supplemented with 10% (v/v) calf serum (Gibco, Thermo Fisher, Waltham, MA, USA) and differentiated at 37°C in 5% CO_2_ using 20 ng/mL phorbol 12-myristate 13-acetate (PMA; Sigma, P8139) for 24 h. The murine macrophages cell line Raw 264.7 (ATCC) were cultured in DMEM (Corning) supplemented with 10% (v/v) calf serum (Gibco, Thermo Fisher) as well. Fresh pre-heated culture medium was added before an overnight resting period.

Peripheral blood mononuclear cells (PBMCs) were isolated from healthy donated whole blood using Lymphoprep (07851, Stemcell, Canada), then centrifuge 2000 rpm for 20 min. Isolated PBMC were washed with PBS and resuspended by RPMI-1640 which supplemented 50ng/ml recombinant human macrophage colony-stimulating factor (M-CSF) (AF-300-25-500, PeproTech, Cranbury, NJ, USA), then cultured at 37°C in 5% CO_2_ for 5 days to differentiate into human monocyte derived macrophages (hMDMs) before infection.

Bafilomycin A (HY-100558, MCE, Monmouth Junction, NJ, USA) was diluted in dimethyl sulfoxide (DMSO, Sigma-Aldrich, Merck) to 100 μM stock solution and diluted to a 20 nM working solution. Chloroquine (HY-17589A, MCE) was diluted in DMSO to 50 mM stock solution and diluted to a 5 μM working solution. ETC inhibitors including: Oligomycin (HY-N6782, MCE) was diluted in DMSO to 10 μM stock solution and diluted to a 10 nM working solution. Prericidin A (MZ9233, maokangbio, Shanghai, China) was diluted in DMSO to 1 mM stock solution and diluted to a 10 nM working solution. Rotenone (HY-B1756, MCE) was diluted in DMSO to 1 mM stock solution and diluted to a 1 μM working solution. Antimycin A (MS0070, maokangbio) was diluted in DMSO to 2.5 mg/ml stock solution and diluted to a 100 nM working solution.

### Human sample

All human samples used in the present study was approved by the Ethics Committees of the Shenzhen Third Hospital (Shenzhen, China). Written informed consent was provided by all study participants.

### RNA interference

THP-1 macrophages were seeded at a density of 4 × 10^5^ cells per well into 12-well culture plates. After 24 h PMA-differentiation, the THP-1 macrophages were transfected with *MFN1* siRNA (RiboBio, Guangzhou, China) and scrambled siRNA (RiboBio) used as a negative control. The sequence of siRNA targeting *MFN1* is 5’-GCACACTATCAGAGCTAAA-3’. Transfection was operated by using Lipofectamine RNAiMAX (ThermoFisher) according to the manufacturer’s protocol. Fresh media was added after 6 h transfection, then the cells were rested for 48 h.

### RNA preparation, RT-PCR, and real-time PCR

Total cellular RNA was extracted using Total RNA Kit (R6634-02, Omega Bio-tek, Norcross, GA, USA), and the isolated RNA was reverse-transcribed into cDNA with HiScript II Q RT SuperMix (R223-01, Vazyme, Nanjing, China) according to the manufacturer’s instructions. The cDNA was mixed with 2 × SYBR Green qPCR Master Mix (B21202, Bimake, Houston, TX, USA), and the real-time PCR reaction was performed on the 7500 Real-Time PCR System (ABI, ThermoFisher). The primers used for the real-time PCR are listed in Supplementary Table 1.

### Western blot analysis

Differentiated THP-1 macrophages, which were transfected with siRNA and infected with Mtb for 24 and 48 h, were lysed in SDS lysis buffer and subjected to western blot (WB) assay. The protein samples were loaded and separated using 10% or 15% sodium dodecyl sulfatepolyacrylamide gel electrophoresis (SDS-PAGE), then transferred onto a polyvinylidene difluoride membrane (Immobilon-P). The membrane was then blocked with 5% bovine serum albumin (BSA) in 0.1% Tween 20/PBS at room temperature for 2 h and incubated with primary antibody at 4°C overnight. Blots were visualized on Minichemi Chemiluminescence Imaging System (SageCreation, Beijing, China) by adding SuperSignal West Pico PLUS Chemiluminescence Substrate (ThermoFisher). The primary antibodies were rabbit monoclonal anti-MFN1 (D6E2S, CST, Danvers, MA, USA, 1:1000); ribbit polyclonal anti-MFN1 (A9880, ABclonal, Woburn, MA, USA, 1:1000); ribbit monoclonal anti-MFN2 (A19678, ABclonal, 1:1000); ribbit polyclonal anti-OPA1 (A9833, ABclonal, 1:1000); ribbit polyclonal anti-DRP1 (A16661, ABclonal, 1:1000); anti-actin (ab179467, Abcam, Cambridge, UK, 1:5000); ribbit monoclonal anti-LC3B (A19665, ABclonal, 1:1000); ribbit monoclonal anti-ATG5 (D5F5U, CST, 1:1000). The secondary antibodies were goat polyclonal anti-rabbit IgG (HRP) (ab6721, Abcam, 1:5000) and goat polyclonal anti-mouse IgG (HRP) (ab6789, Abcam, 1:5000).

### CFU assay

For the CFU count, 5 × 10^5^ differentiated THP-1 macrophages were seeded into each well of a 12-well plate and infected with mycobacteria, as described previously. After 6 h of infection, extracellular bacteria were removed by washing twice with sterile PBS. Fresh pre-heated culture medium was added, and the cells were rested for 72 h. Extracellular bacteria were removed by washing twice with sterile PBS. The cells were lysed in 0.1% SDS gradient dilution in sterile PBS, then plated onto 7H10 agar plates. The plates were incubated at 37°C with 5% CO_2_ for 2-3 weeks, and the colonies were enumerated.

### Immunofluorescence

Differentiated THP-1 macrophages were grown on coverslips at a low density (2 × 10^5^ per well), followed by transfection with siRNA and infection with Mtb for 48 h. The cells were fixed for 15 min in 4% paraformaldehyde (Biosharp, South Jordan, UT, USA) at 37°C, permeabilized for 10 min in 0.03% TritonX-100, and blocked for 30 min with 3% BSA in 0.1% Tween 20/PBS. The cells were incubated with mouse monoclonal anti-TOMM20 antibody (ab56783, Abcam, 1:1000) overnight at 4°C. On the second day, the cells were equilibrated to room temperature for 1 h, washed with PBS, incubated for 1 h with secondary antibody Alexa Fluor 555 goat anti-mouse IgG (H+L) (A21422, ThermoFisher), washed with PBS, incubated with Hoechst 33342, washed with PBS, and treated with antifade solution. The cells were visualized under a 60 × oil objective of a confocal microscope (Nikon A1R). The mitochondrial morphology was scored in at least 200 cells per group by using two different categories of morphology (fragmented and tubular). Fragmented group was defined as cells with majority (>50%) fragmented mitochondria, tubular group represent cells over 80% elongated mitochondria (41, 42). Scoring was preformed blind.

JC-1 (C2006, Beyotime, Shanghai, China) assay which to determine MMP was also carried on confocal microscope according to the manufacturer’s instructions.

### MMP assays, mROS assays and Flow cytometry

For analysis of MMP, infected or uninfected THP-1 macrophages were stained with 100 nM MitoTracker Red CMXRos (M7512, ThermoFisher) or 1 μM rhodamine 123 (C2007, Beyotime) in culture medium for 30 min at 37°C After washing twice with PBS, the cells were processed for flow cytometry. For analysis of mROS release level, infected or uninfected THP-1 macrophages were stained with 10 *μM* MitoSOX Red mitochondrial superoxide indicator (M36008, Invitrogen, Thermofisher). Flow cytometry was carried on BD FACSAriaII cell sorter (BD biosciences) and analysis using FlowJo software.

### ATP

The intracellular ATP levels of THP-1 macrophages was evaluated using the Enhanced ATP Assay Kit (S0027, Beyotime) according to the manufacturer’s instructions.

### OCR assay

The OCR and ECAR of THP-1 macrophages were determined using the Seahorse XFe24 Extracellular Flux Analyzer (Agilent Technologies, Santa Clara, CA, USA). Cells were seeded at 8 × 10^4^ per well, differentiated with PMA for 24 h, and infected as previously described. We used the Seahorse XF Cell Mito Stress Test Kit (103015-100, Seahorse Bioscience) for the OCR assay. Electron transport chain inhibitors were added every 20 min from the beginning in the following order: oligomycin (1.5 μM), FCCP (1 μM), followed by a combination of rotenone (0.5 μM) and antimycin A (0.5 μM). For the ECAR assay, the Seahorse XF Glycolysis Stress Test kit (103020-100, Seahorse Bioscience) was used. All readings were normalized to the protein level of each well using the BCA Protein Assay Kit (P0011, Beyotime).

## Statistical analysis

Statistical analysis was performed using GraphPad Prism 6 software (GraphPad Software, Inc.). A two-tailed unpaired Student’s t-test was used to evaluate the significance of single parameters between two groups. One-way analysis of variance (ANOVA) with multiple comparisons was used to analyze more than two groups. Data were expressed as mean ± SEM (n = 3). Significance is presented as *, p<0.05; **, p<0.01; ***, p<0.001; ****, p<0.0001; “ns”, not significant.

## Acknowledgments

We thank Jessica Tamanini (Shenzhen University and ET editing) for English editing before submission. We also thank Hongbo Shen (Institute Pasteur of Shanghai, Shanghai, China) for gifting mRFP-GFP-LC3B THP-1 cells and kindly help in detecting and analyzing autophagic flux.

This project was supported by the Natural Science Foundation of China (82072252 and 91942315) and the Guangdong Provincial Key Laboratory of Regional Immunity and Diseases (2019B030301009).

The funders had no role in the study design, data collection and analysis, decision to publish, or preparation of the manuscript.

---

**Supplementary Figure 1.**
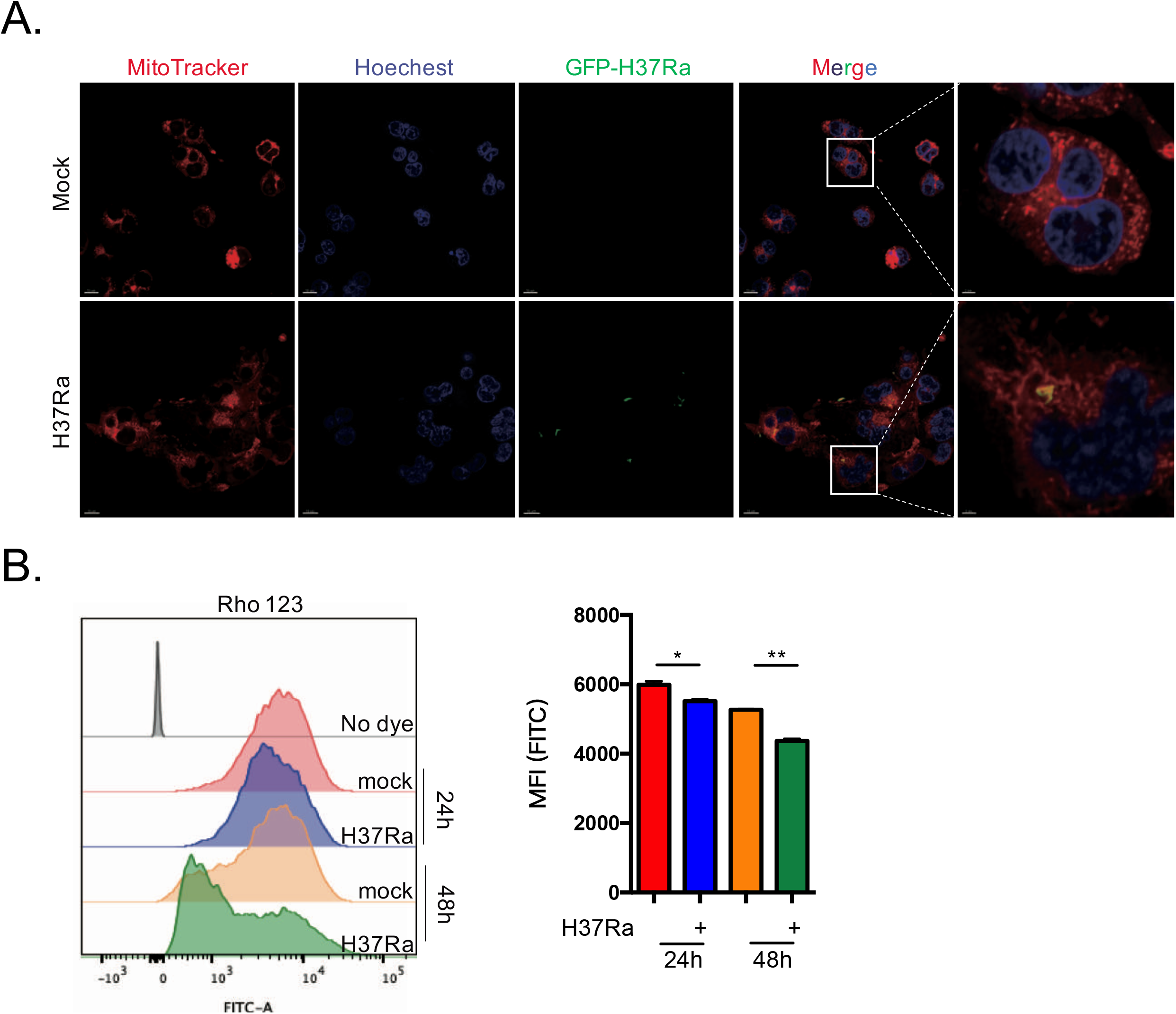
Mtb increases mitochondrial fusion and mitochondrial membrane potential in infected THP-1 macrophages. A. Confocal microscopy images showing mitochondrial fusion in H37Ra- (MOI = 10:1) infected THP-1 macrophages for 48 h. The cells were stained with MitoTracker Red CMXRos probes (100 nM). Red fluorescence, mitochondria; green fluorescence, H37Ra; blue fluorescence, Hoechst 33342-labeled nuclei. (Scale bars, left 10μm, right 3μm) B. Mitochondrial membrane potential (MMP) was detected using rhodamine (Rho) 123 (1 μM) for the H37Ra (MOI = 10: 1) infected THP-1 macrophages. Mean fluorescence intensity (MFI) of MMP was analyzed by flow cytometry. The data represent mean ± SEM (n = 3); *, p < 0.05; **, p < 0.01.

**Supplementary Figure 2.**
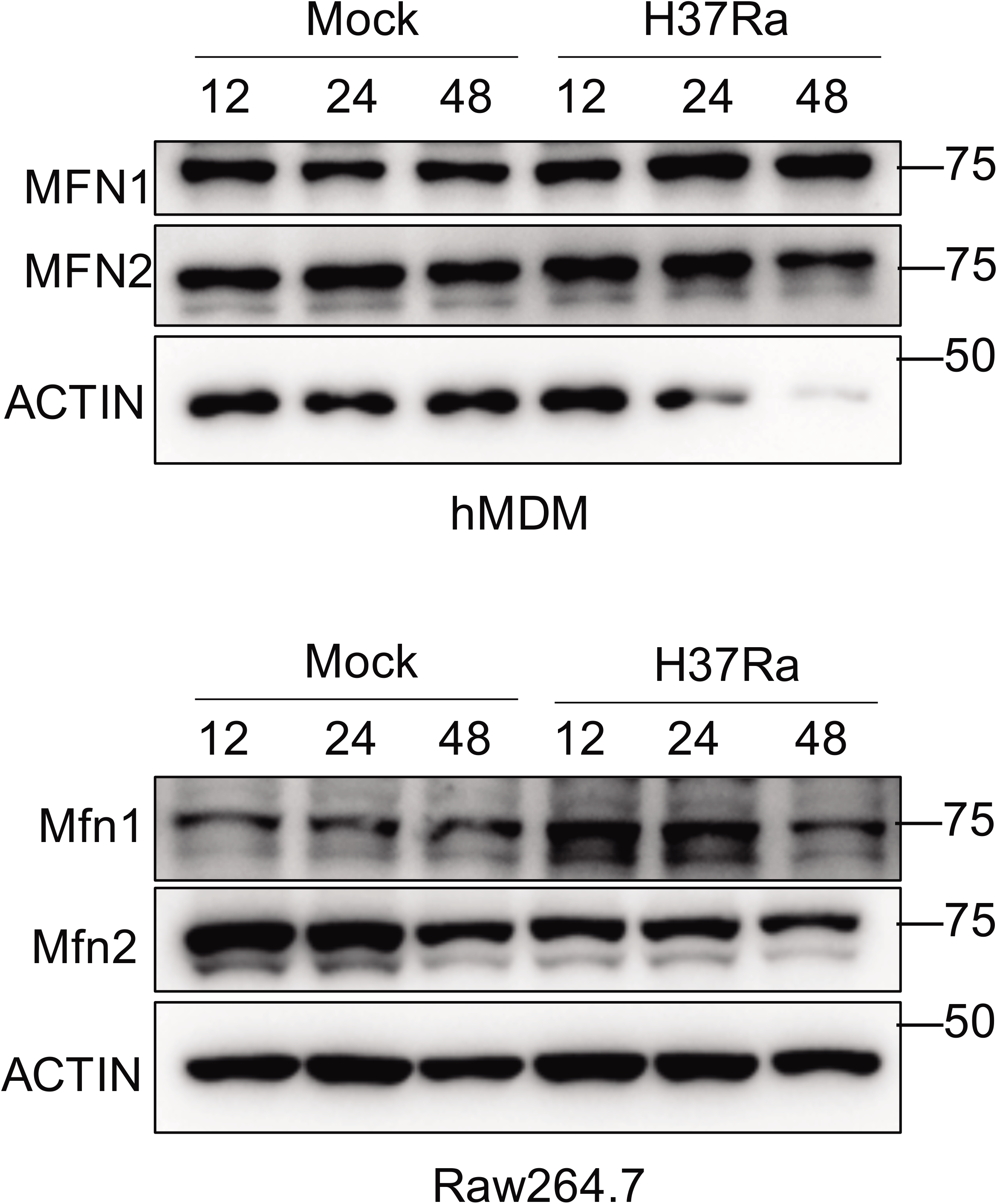
Expression of mitofusin1 and mitofusin 2 in hMDMs and Raw264.7. Cell lysates were harvested from uninfected (mock) and 12, 24, and 48 h H37Ra (MOI = 10: 1) infected human monocyte-derived macrophages (hMDMs) and Raw264.7. Western blot analysis of MFN1 and MFN2; β-actin was used as the loading control.

**Supplementary Figure 3.**
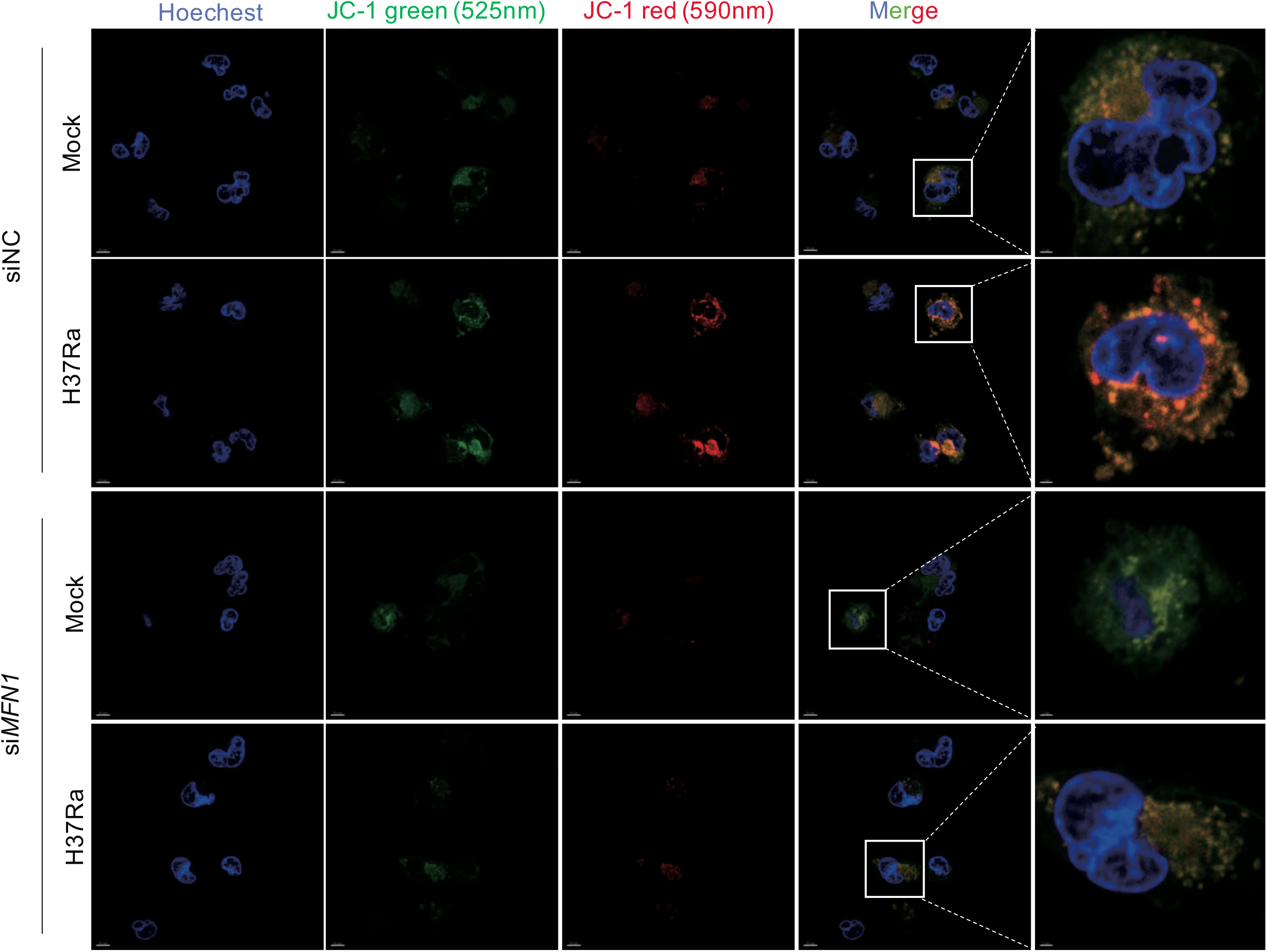
*siMFN1* leads to decrease in mitochondrial membrane potential. Confocal microscopy images showing MMP (JC-1 staining) in uninfected or H37Ra (MOI = 10: 1) infected THP-1 macrophages at 48 hpi in the present or absence of *MFN1* siRNA compared with negative control siRNA (siNC). Images shown increased MMP in H37Ra infected siNC group macrophages and decreased MMP upon *MFN1*-siRNA treatment. JC-1 monomers which show green florescence fused into mitochondrial and form red florescence aggregates depends on higher membrane potential. (Scale bars, left 10μm, right 2 μm).

**Supplementary Figure 4.**
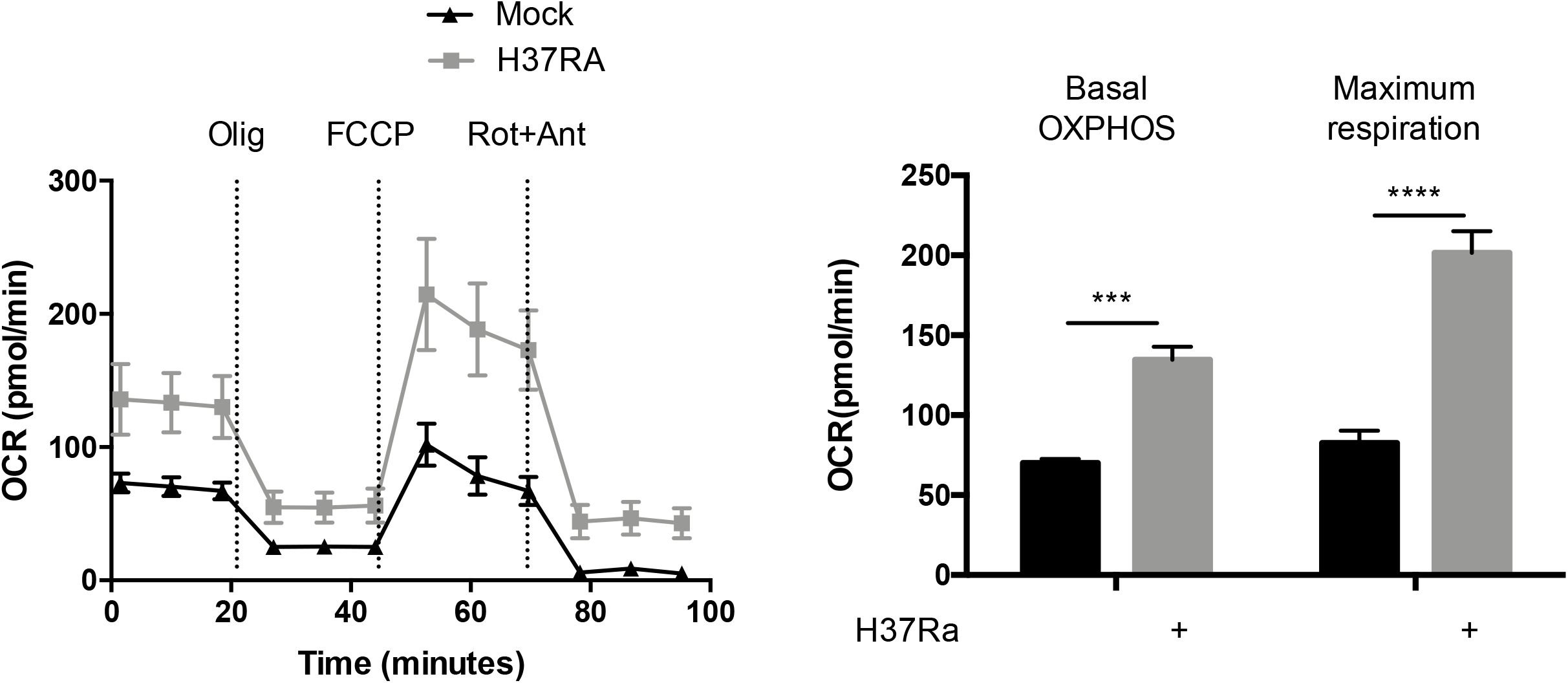
Elevated OXPHOS in THP-1 macrophages 24 h after Mtb-infection. Mito Stress assay showing increased OCR of THP-1 macrophages after infection by H37Ra (MOI = 10:1) for 24 h. The data represent mean ± SEM (n = 3); ***, p<0.001; ****, p<0.0001.

**Supplementary Figure 5.**
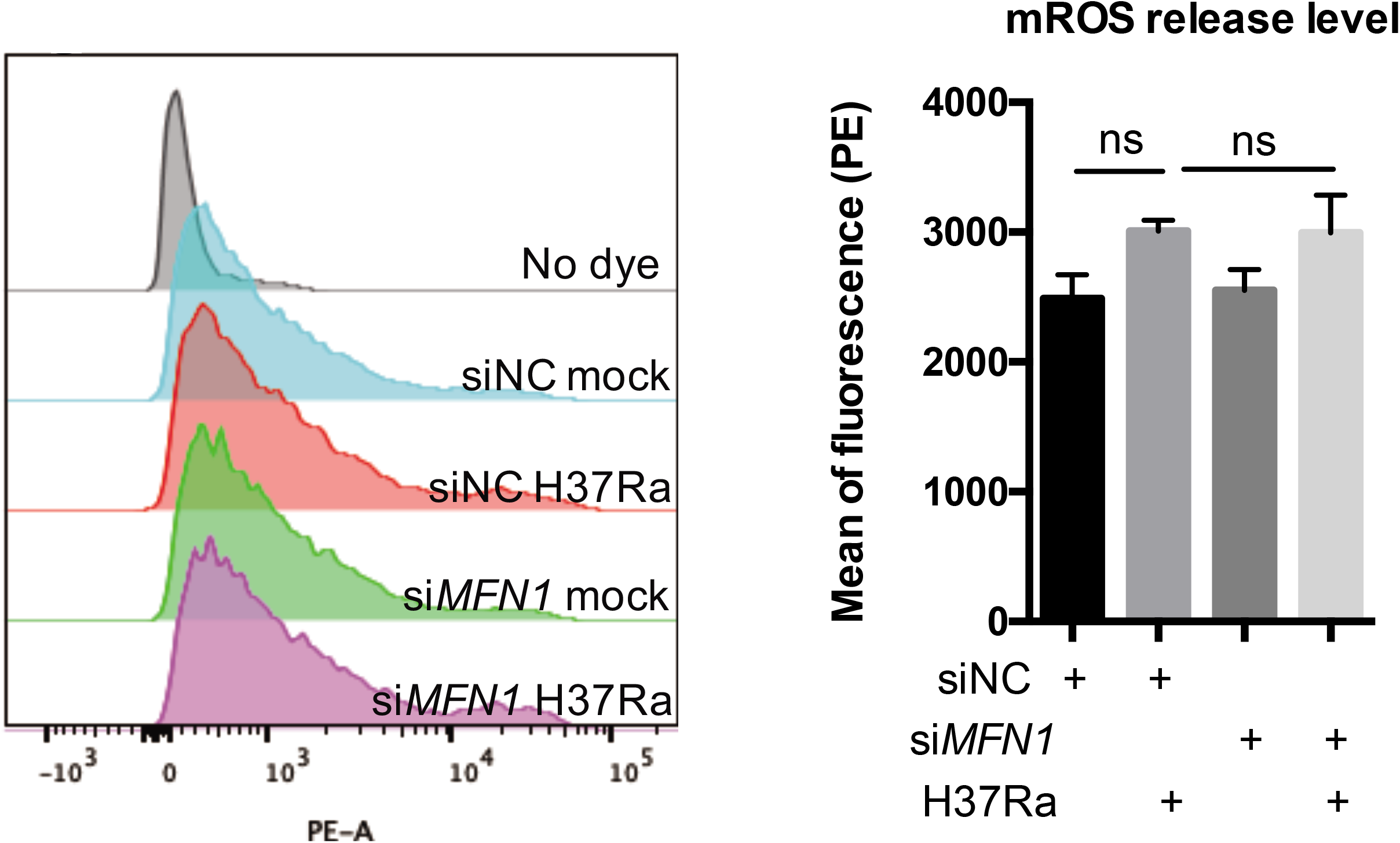
There was no significant difference in mROS release between si*MFN1* group and control group. Levels of mROS released by uninfected or H37Ra (MOI = 10: 1) infected-THP-1 macrophages which treated with or without *MFN1* siRNA compared with negative control siRNA (siNC) were detected using MitoSOX probes (10 uM) and analyzed by flow cytometry at 6 hpi. Histograms indicated mean fluorescence intensity (MFI) of mROS release level analyzed by flow cytometry. The data represent the means ± SEM (n = 3); “ns”, not significant.

**Supplementary Figure 6.**
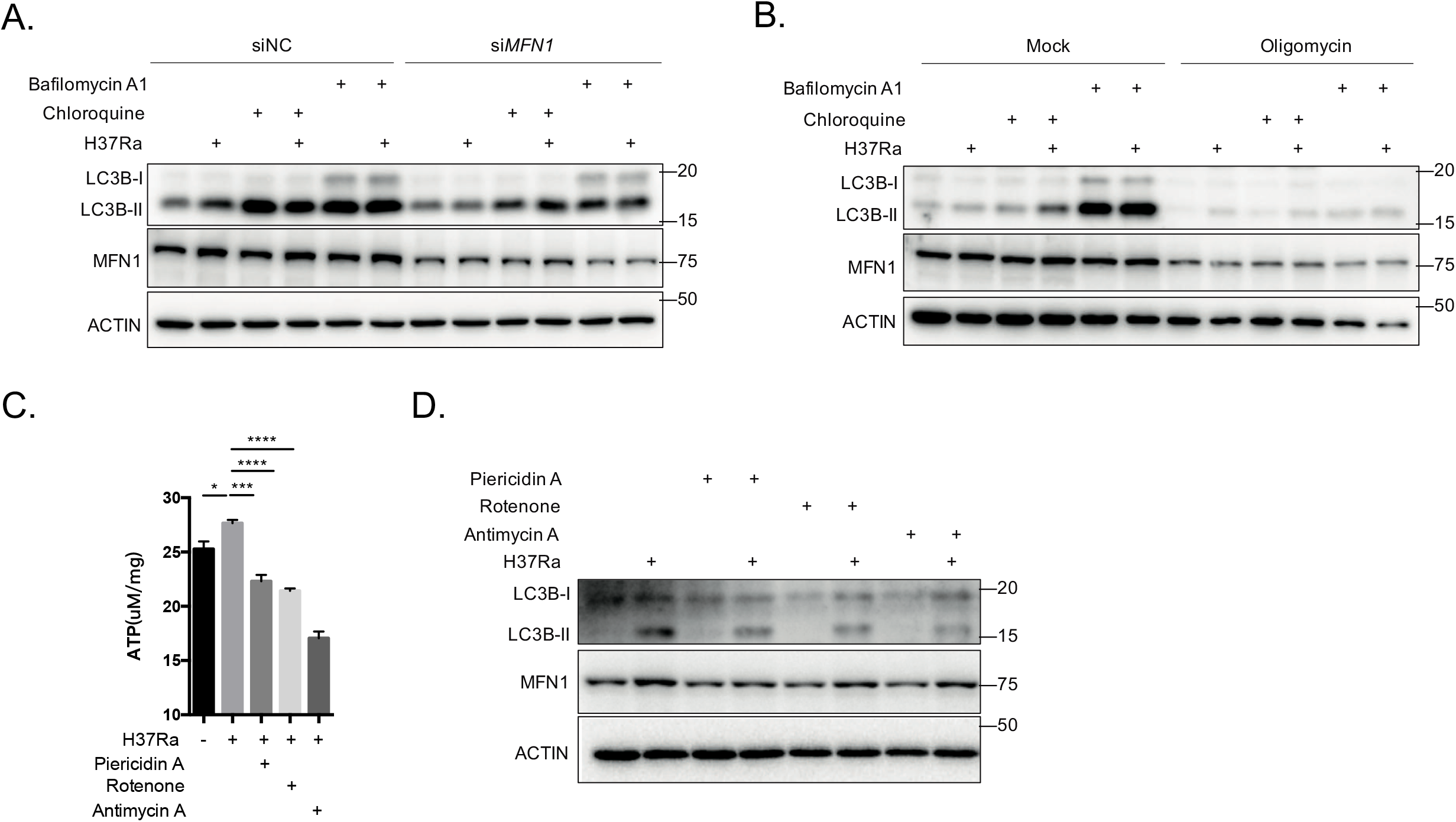
Autophagy was inhibited by si*MFN1* and ETCi. A-B. Cell lysates were harvested from uninfected (mock) and H37Ra (MOI = 10: 1) infected THP-1 macrophages treated with or without *MFN1* siRNA compared with negative control siRNA (siNC) at 24 hpi (A) and oligomycin (B). Chloroquine (5 M) and bafilomycin A1 (20 nM) were used as autophagy inhibitors which inhibit autophagosome-lysosome fusion and lead accumulation of LC3B-II. These results demonstrate si*MFN1* inhibited the initiation of the autophagy process. Western blot analysis of MFN1 and LC3B; “-actin was used as the loading control. C-D. Electron transport chain inhibitors (ETCi) including piericidin A (10 nM) (inhibitor of complex I) and rotenone (1 μM) (inhibitor of complex I) and antimycin A (100 μM) (inhibitor of complex III) were added to the uninfected or H37Ra (MOI = 10: 1) infected-THP-1 macrophages. Total intracellular ATP was measured at 24 hpi using luminescent assay (C). Cell lysates were harvested 24 hpi for western bolt analysis (D). The data represent the means ± SEM (n = 3); *, p < 0.05; ***, p < 0.001, ****, p < 0.0001.

**Supplementary Table 1.**
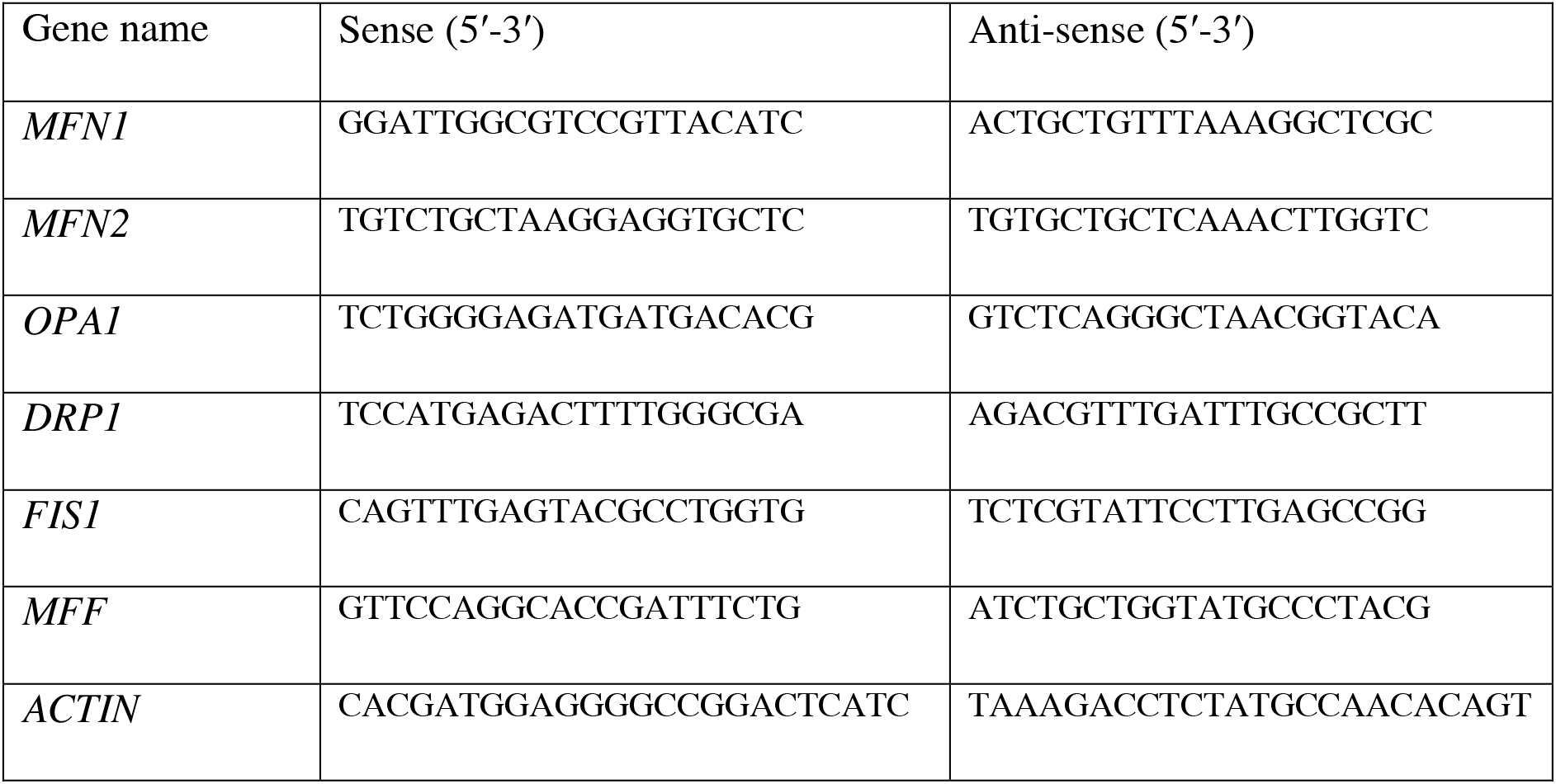
Real-time PCR primer sequences for mRNA expression in the study.

